# Competitive interactions between motor and episodic memory systems during interleaved learning

**DOI:** 10.1101/2020.01.10.901207

**Authors:** Sungshin Kim

## Abstract

Distinct motor and episodic memory systems are widely thought to compete during memory consolidation and retrieval, yet the nature of their interactions during learning is less clear. Motor learning is thought to depend on contributions from both systems, with the episodic system supporting rapid updating and the motor system supporting gradual tuning of responses by feedback. However, this competition has been identified when both systems are engaged in learning the same material (motor information), and so competition might be emphasized. We tested whether such competition also occurs when learning involved separate episodic-memory and motor information presented distinctly but yet in close temporal proximity. We measured behavioral and brain-activity correlates of motor-episodic competition during learning using a novel task with interleaved motor-adaptation and episodic-learning demands. Despite unrelated motor versus episodic information and temporal segregation, motor learning interfered with episodic learning and episodic learning interfered with motor learning. This reciprocal competition was tightly coupled to corresponding reductions of fMRI activity in motor versus episodic learning systems. These findings suggest that distinct motor and episodic learning systems compete even when they are engaged by system-specific demands in close temporal proximity during memory formation.

## 1. Introduction

Distinct neural systems supporting motor versus episodic memory ^1^ are thought to operate simultaneously during motor learning, such as when one must adapt motor output to accommodate externally applied perturbations ^2–6^. Episodic memory for goals, strategies, and feedback may dominate the earlier stage of learning, with gradual evolution of implicit motor memory that dominates later stages of learning and produces automaticity ^7–11^. Thus, independent learning processes with distinct timescales operate simultaneously to achieve common learning goals, such as improving motor acuity, accuracy, and response times. Thus, interference between motor and episodic learning can also occur via competition while achieving the same goal for episodic and motor memory systems, probably due to limited memory resource shared by the systems. For instance, in the consolidation period after motor learning, presentation of word-list material that is learned episodically interferes with subsequent expression of the previous motor learning, suggesting episodic-motor competition during consolidation that may arise from consolidation bandwidth limits ^12–16^. However, these studies retrospectively inferred interaction between the memory systems based on performance of tasks after consolidation period without any direct demonstration of underlying neural signatures ^12,13,16^ or pre-assumed that specific brain regions (e.g., DLPFC, M1) are associated with separate memory processing ^14,15^. Moreover, previous studies have shown competition either during learning for the common task goals or after learning for the separate system-specific task goals. However, it has been unclear whether the interactions would be competitive or cooperative during learning for the system-specific task goals. To fill this gap, we aimed to investigate direct evidences of the interaction between episodic and motor systems during learning by analyzing trial-by-trial performance of distinct episodic versus motor tasks.

Here, we developed a novel fMRI task in which subjects performed visuomotor adaptation, which is a standard type of motor learning ^3–5,17,18^, interleaved with object-location association learning, which is a standard type of hippocampal-dependent episodic learning ^19–22^ (Figure 1). Visual feedback on which motor learning is dependent was either presented or hidden and trial-level motor error was used to quantify the amount of motor learning that occurred from one trial to the next. Additionally, we also designed a control motor task with feedback but without visuomotor adaptation to rule out possible effects of non-specific factors except for memory interference by the visual feedback *per se*, e.g., attention. Episodic memory testing after the learning phase was used to quantify the relative success of episodic learning on each trial with measuring recognition with the level of confidence and recollection of associated locations. This allowed us to quantify the degree to which motor and episodic learning occurred on a trial-by-trial basis. According to the hypothesis of competitive interaction between the memory systems, we predicted that motor learning would harm episodic learning and that the magnitude of this episodic memory disruption would scale with the magnitude of trial-by-trial motor learning. Likewise, we predicted that successful episodic learning would negatively affect the success of motor learning in temporally proximal trials.

**Fig. 1.**
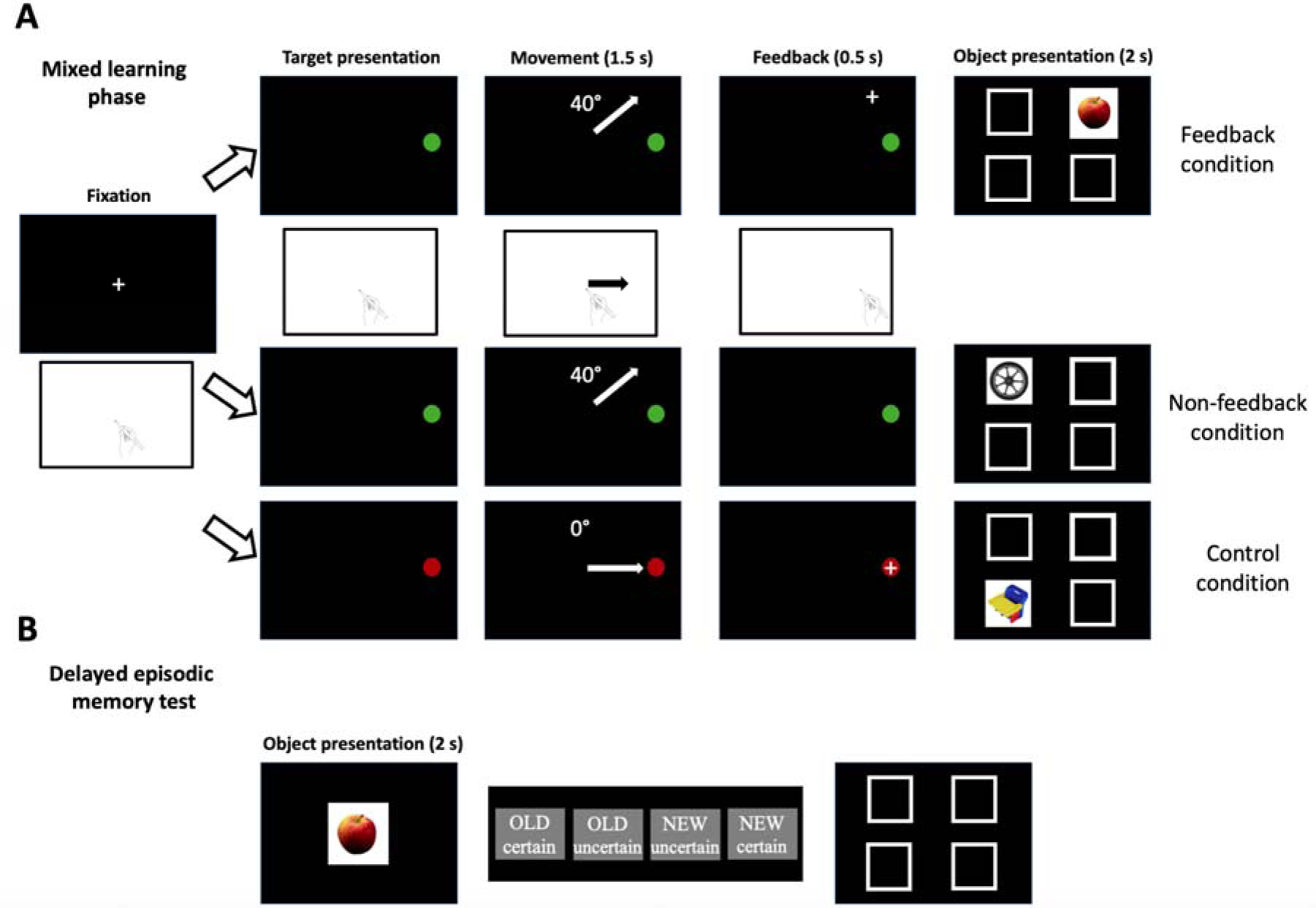
Experiment design. (**A**) For each trial in the interleaved learning phase, a target appeared on the right side of the screen and subjects attempted to reach the target using an fMRI-compatible pen on a tablet within 1.5 s (maximum movement time). Target colors cued different visuomotor rotations, 0°, +40°, and −40° (−40 degrees not illustrated here). In the feedback and control conditions, a cursor position was provided as feedback for 0.5 s after the movement. Immediately after feedback, a trial-unique object appeared in the one of four locations. (**B**) In the delayed episodic memory test, we assessed object recognition memory and recollection memory of the associated location.

To identify systems-level interactions supporting the predicted bidirectional interference of motor and episodic learning, we fit trial-by-trial measures of motor learning and episodic learning to fMRI activity. We predicted that activity of a motor-learning network including prefronto-parieto-cerebellar areas would positively correlate with the success of motor learning and would negatively correlate with the success of episodic learning. Likewise, we predicted that the negative influence of motor on episodic learning would correlate with reduced activity in an episodic-memory network including hippocampal and medial-prefrontal areas. This predicted pattern of findings would provide evidence for general competition between distinct motor and episodic memory systems; that is, competition even when learning involves distinct system-specific information presented in relatively close temporal proximity.

## 2. Methods

### 2.1. Participants

Behavioral and fMRI data were collected from twenty-five right-handed and neurologically healthy subjects (13 females, mean age = 25.7 years, age range = 19 - 35 years). Handedness was assessed by a modified version of the Edinburgh Handedness Inventory (score ≥ 50, for right-handedness ^23^). All participants had normal or corrected-normal vision and were eligible for the experiment based on standard MRI safety screening. They gave written informed consent in accordance with the Declaration of Helsinki and were remunerated for their participation. The experimental protocol received approval from Northwestern University Institutional Review Board. Two subjects were excluded for no learning of motor tasks (one falling asleep in the scanner; see results for detailed exclusion criteria). Therefore, twenty-three subjects were included in data analysis (13 females, mean age = 25.7 years, age range = 19 - 35 years)

### 2.2. Task procedures and experimental design

We designed an interleaved task-based fMRI experiment with motor learning and object-location association memory demands. On the beginning of each trial, a round target of 1 degree in visual angle appeared on the right of the screen, 5.8 degrees of visual angle away from the center. Subjects were instructed to manipulate an MRI-compatible tablet pen (Hybridmojo LLC, CA, USA) to move the cursor to the target within 1.5 s after the target onset (maximum movement time). Timed-out trials were considered invalid and excluded from data analysis (0.92 % of all trials). There were five different experiment conditions; a control condition with visual feedback of the cursor movement, two visuomotor task conditions, in which the cursor movement was rotated 40° (clockwise, Task 1) and −40° (counter-clockwise, Task 2) with or without the visual feedback (Figure 2). Three different target colors, yellow, blue, green, were used to differentiate the control task and two visuomotor tasks.

**Fig. 2.**
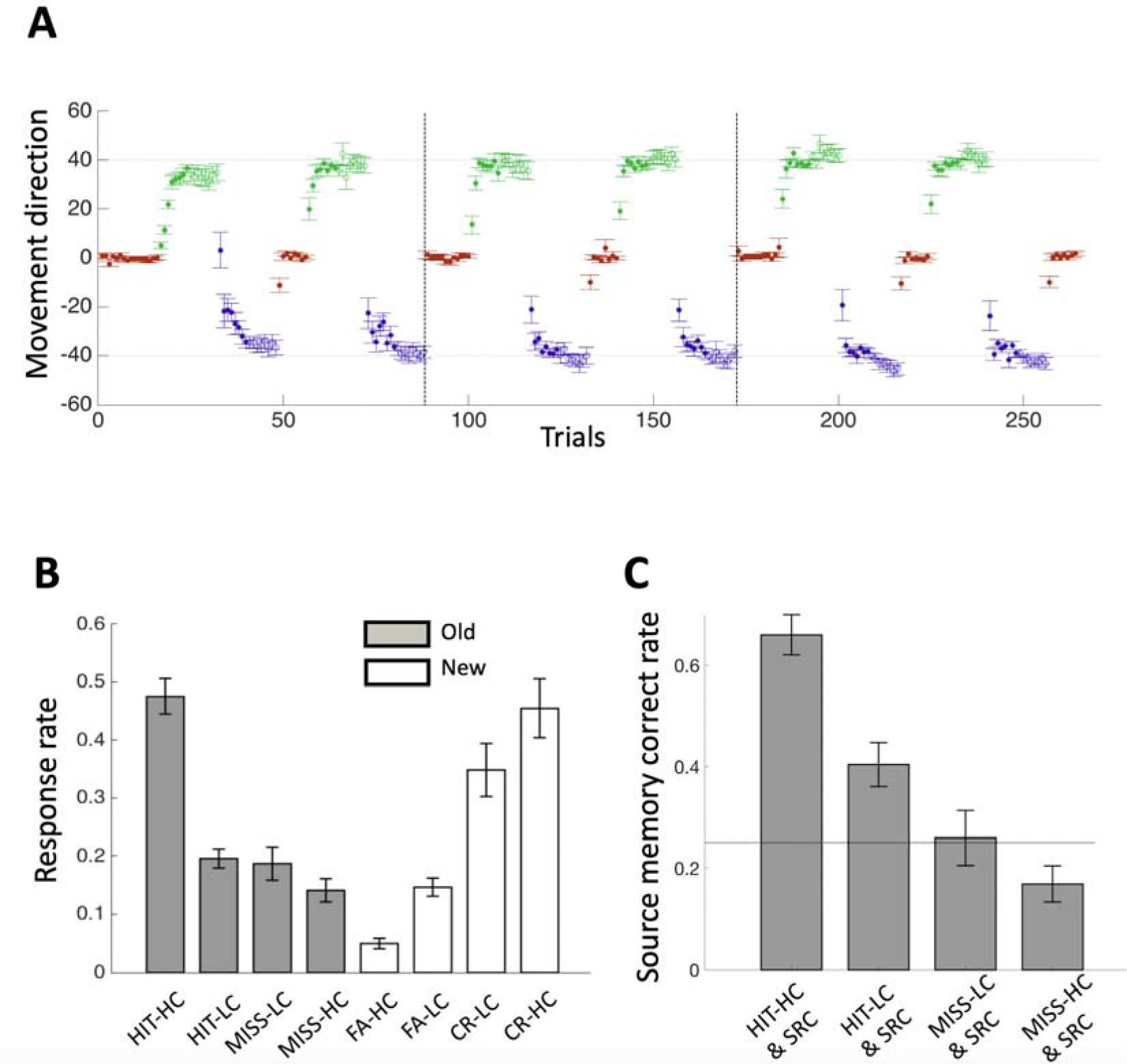
Motor and episodic learning. (**A**) Red, green, and blue circles indicate movement directions in Control (0° rotation), +40° Feedback, and −40° Feedback conditions, respectively, averaged across participants. The open circles indicate the non-feedback condition. (**B**) Responses rates for episodic recognition memory were averaged separately for old versus new objects and for each confidence level (HC: high confidence; LC: low confidence) and each response category (HIT, MISS for old objects, FA: false alarm, CR: correct rejection for new objects). (**C**) Correct response rates for source memory are shown averaged given the corresponding recognition response. The dotted line indicates the chance performance level.

For the two visuomotor tasks with feedback, a cross-shaped white cursor (0.83 degree in visual angle) appeared as a current position (or rotated in visuomotor learning blocks) of the tip of the pen while movement, relatively from the center of the screen (shown as the initial position regardless of actual position in the tablet). Once the cursor crossed over 5.8 degrees of visual angle from the center, the cursor was fixed at the crossing point. The cursor was presented for additional 0.5 s after the maximum movement time. For the two visuomotor tasks with non-feedback, a cursor was not presented but subjects were instructed to move the pen to the target as if there was the cursor. With color codes of round targets, subjects could recognize a current task even without cursor feedback. Immediately after the round target disappeared, a trial-unique object (3.3 x 3.3 degrees of visual angle ^24^) appeared in one of four locations with pseudo-random order (permuting the four locations, Figure 1) for two seconds. Subjects were instructed to remember objects with their appeared locations and move back to the home position. Object images were taken from stimulus sets developed for research purposes, which were made publicly available with no copyright protection via the internet (Computational Visual Cognition Lab; http://cvcl.mit.edu/MM; Cambridge, MA). Inter-stimulus-intervals (ISI: time during a fixation cross is presented between trials) were randomly generated from 2 s to 12 s from an exponential distribution, in 2 s increment.

The fMRI experiment was divided in three runs, lasting 700, 668, and 730 s, respectively with a short break (~30 s) between runs. Each experimental block consisting of eight trials and was presented according to schedules such as C11’22’C11’22’ (12 subjects) or C22’11’C22’11’ (11 subjects) per run, where C, 1, and 2 indicates a block of the Control, Task 1, and Task 2 and the dash mark indicates non-feedback block of the equivalent task. Thus, there were 30 experimental blocks in three runs, 6 blocks for each condition (Control, Tasks 1 and 2 with or without feedback). Possible confounding effects due to schedule and target color were eliminated by counter-balancing two schedules and three color-codes across subjects. To avoid primacy and recency effects ^25^, eight control task trials were added to the beginning and the end of the experiment, and four control task trials were added to the beginning of second of the third runs, constituting a total of 264 trials. These trials were not included main analyses except in Figure 2A showing entire course of learning. Subjects practiced a familiarization session of ~100 trials of the control task before the experiment. Learning two opposing visuomotor rotations while performing the associative memory task takes hundreds of trials, thereby providing enough statistical power in fMRI analysis as well as making more likely interaction between motor memory and episodic memory.

In a delayed (~10 minutes) memory test session outside of a scanner, 528 objects were presented, half of which were old (presented in the fMRI experiment) and the other half new, in randomized order. On each trial, the object was presented in the center of computer screen for two seconds and a prompt appeared asking subjects to classify the object as old or new, each with two confidence levels, “certain” or “uncertain”. For old objects only, subjects were then asked to click one of four appeared boxes corresponding remembered location or click anywhere outside of four boxes if they do not remember. There was no time limit for these responses and two-second interval with the fixation-cross separated the response period from the next trial. The test session typically lasted approximately 45 minutes depending on responses of subjects.

Stimuli were presented on a computer screen and reflected onto a mirror mounted to the head coil. Subjects laid in a supine position in the scanner and had a tablet fixed on their lap so that they could reach all the corners of the tablet comfortably with their right hand.

### 2.3. Behavioral data analysis

For each of 264 trials, we calculated the average and variability of movement direction across subjects (Figure 2). The angular error was calculated as a size of deviation between the target direction and the final cursor direction from the center of the screen. Trial-by-trial movement direction were averaged across subjects and displayed with error bars (SEM) to show overall performance of motor learning.

Recognition memory performance as fraction of later-hit trials was calculated within each of 30 experimental blocks except for additional blocks of primacy and recency effects (Figure 2). Source-recollection memory performance was calculated as the fraction of correct responses out of valid responses (i.e., click one of four response boxes). Note that, for five subjects, there were no valid trials used to calculate source memory scores when they failed to recognize objects because they responded as “don’t remember” for all the trials by clicking outside of four response boxes. The recognition memory performance was highly correlated with source-memory performance across subjects and thus we did not perform a separate analysis for the source-memory recollection.

The recognition memory performance was compared among three conditions, the control, feedback, and non-feedback across six experimental blocks (two for each run) and post-hoc paired t-tests were performed. We hypothesized the recognition memory performance in the feedback condition would be lower than those in the control and the non-feedback condition due to interference of motor learning (i.e., motor memory update) with episodic learning. We also estimated motor learning within 30 experiment blocks and related with the recognition memory performance. For each of 23 subjects, movement directions within each block were fitted by an exponential function with three free parameters (*A*, *B*, *C*) such that *x*(*t*) = *A* • exp(−*B*(*t* − 1)) + *C*, where *x*(*t*) is a movement direction at trial *t*, (*t* = 1, 2,…,8). For robust estimation of learning with few trials, we excluded missed trials and trials with large (>20°) overshoot (less than 3.5%). Learning within a block was estimated as the amount of movement change on the fitted curve from the initial to the final trial. Then, the estimated amounts of learning within a block were averaged across subjects and correlated with averaged recognition memory performance.

We also investigated on the interference of the opposite direction, that is, from episodic learning to motor learning. For this, the 240 studied objects except for those used for primacy and recency effects (24 objects) were back-sorted to be categorized into four different levels of episodic memory formation success according to their responses in a delayed test session; later-miss (MISS: Level 1), later-hit with low-confidence (HIT-LC: Level 2), later-hit with high confidence (HIT-HC: Level 3), later-hit with high-confidence and source-correct (HIT-HC & SC: Level 4). Then, we compared the trial-by-trial motor learning associated with the object, which was defined as the change of errors in motor learning task from the current to the next trials. Here, we hypothesized higher episodic memory formation success for a given object would more interfere with motor learning.

### 2.4. MRI data collection and preprocessing

MRI data were acquired using a 64-channel head/neck coil on a 3-tesla Siemens TIM Prisma whole-body scanner at Northwestern University Center for Translational Imaging. A high-resolution T1-weighted structural 3D MP-RAGE was acquired before the task to provide anatomical location (voxel size: 1mm^3^; field of view: 256 mm, 151 sagittal slices). Whole-brain functional images were acquired during the fMRI experiment, 350, 334, and 365 scans for three runs, respectively. Scanning parameters were 2000-ms repetition time (TR), 20-ms echo time (TE), 210 mm field of view (FOV), 80° flip angle (FA), 1.7×1.7×1.7 mm isotropic voxels, and multi band factor of 2. Processing of fMRI data used freely available AFNI software ^26^. All functional images were first corrected multi-band slice-timing (3dTshift), and then realigned to adjust for motion-related artifacts (3dvolreg). The structural image was skull-stripped (3dSkullStrip) and coregistered with functional images using normalized mutual information as a cost function (align_epi_anat.py, with option, “-nmi”). The realigned images were then spatially normalized with the MNI152 T1 template (@auto_tlrc). All functional images were smoothed using a Gaussian kernel of 4×4×4 mm full width at half maximum (3dmerge) and scaled to mean of 100 for each run (3dcalc).

### 2.5. fMRI data analysis

We performed two regression analyses using parametric regressors on fMRI data to identify activity supporting bidirectional interference shown in behavioral results (see Results) and psychophyisiological interaction (PPI) analyses ^27^ to identify functional coupling between motor and episodic memory networks. First, for the interference from motor learning to episodic learning, we designed a parametric regressor. Specifically, a boxcar function with amplitude modulated by trial-by-trial motor error, onsets and duration encoding 2s-long presentation of object stimuli was convolved with a canonical hemodynamic. Here, we made important assumption that motor learning is proportional to motor error in the visuomotor adaptation task, as has been supported by previous studies ^2–4,13,17,28–31^. For regressors of non-interest, we added boxcar functions encoding blocks of primacy and recency trials, six rigid-body motion regressors and six regressors for each run modeling up to 5th order polynomial trends in the fMRI time series. A beta-value of the error-modulating regressor was estimated via a general linear model incorporating hemodynamic response deconvolution (3dDeconvolve).

From the first regression analysis, we defined two distinct networks, which were constructed by clusters of significantly positive and negative beta-values of the error-modulating regressors. For multiple comparison correction, voxel-wise threshold was set to *p* < 0.001 two-tailed and a Monte Carlo simulation determined 38 contiguous supra-threshold voxels (187 mm^3^) was needed to achieve cluster-wise corrected threshold *p* < 0.05 within the whole-brain group mask (3dttest++ with the option “-Clustsim”) ^32^. Each of positive and negative interaction networks consisting of the significant clusters was overlaid over the MNI template brain.

Then, we sought to test whether the deactivation of the hippocampal-prefrontal memory network negatively interacting with motor learning, is correlated with the extent of the interference with episodic memory formation due to motor learning. For this, the averaged beta-values of the error-modulating regressor in the vmPFC (the most robust cluster identified by the primary fMRI analysis) were correlated with the episodic memory accuracy reduction in feedback blocks compared to following non-feedback blocks across subjects (see Figure 3A).

**Figure 3.**
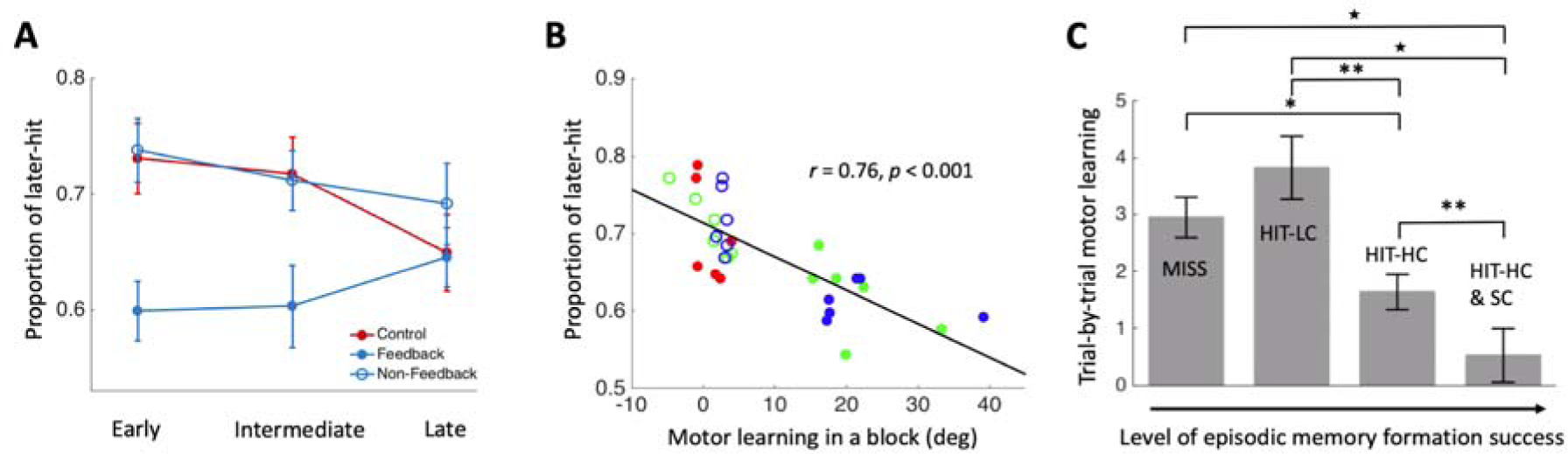
Reciprocal interference between motor and episodic learning. (**A**) Recognition memory performance (blue) is significantly lower in feedback blocks (in which motor learning occurred) than in the non-feedback and the control blocks that were devoid of motor learning and that controlled for various nonspecific factors (see text). (**B**) Episodic and motor learning levels correlated negatively across blocks, with conditions indicated via coloration as in Figures 1 and 2. (**C**) There was less motor learning (improvement from one trial to the next) with increasingly more successful episodic learning. (^⍰^*p* < 0.001, \*\**p* < 0.01, \**p* < 0.5).

Second, for the interference from episodic memory formation to motor learning, we generated four regressors encoding the onsets of motor task feedback only in the trials of the feedback blocks, which were classified depending on a response in a delayed test session; ‘hit with high confidence with source-memory correct’, ‘hit with high confidence with source-memory incorrect’, ‘hit with low confidence’, and ‘miss’. Each regressor consisted of eight tent functions starting 1 s to 15 s after the target onset of motor task with peaks every 2 seconds aligned to TR. Experiment blocks of recency & primacy, control and non-feedback condition were separately modeled as regressors of non-interest. The other regressors of non-interest modeling head motion and polynomial trend were the same as those of the first regression analysis.

In the second regression analysis, we searched neural correlates of the interference for the opposite direction, from episodic memory formation to motor learning. To this end, the beta-values within the positive-interaction network were averaged for each of four regressors of the second regression analysis corresponding to a varying level of episodic memory success. To compare with behavioral results, we calculated the mean of two estimated beta-values of regressors, for “hit with high confidence” regardless of source memory response.

Last, in the PPI analyses, we first generated the physiological regressors as timeseries extracted from each of the three clusters in the hippocampal-prefrontal memory network defined by the first regression analysis, vmPFC and bilateral hippocampus (see Table 1). For the psychological factors, we used three contrasts of hit versus miss, hit with high confidence versus everything else and among three conditions of the motor task, control, feedback, and non-feedback. For each of nine PPI analyses, which are combinations of three physiological regressors from the seed regions and three psychological regressors, we also included regressors encoding the onsets of a target separately for the psychological factors (psychological regressors), as well as a physiological regressor, in addition to the interaction regressor between physiological and psychological factors.

**Table 1.**
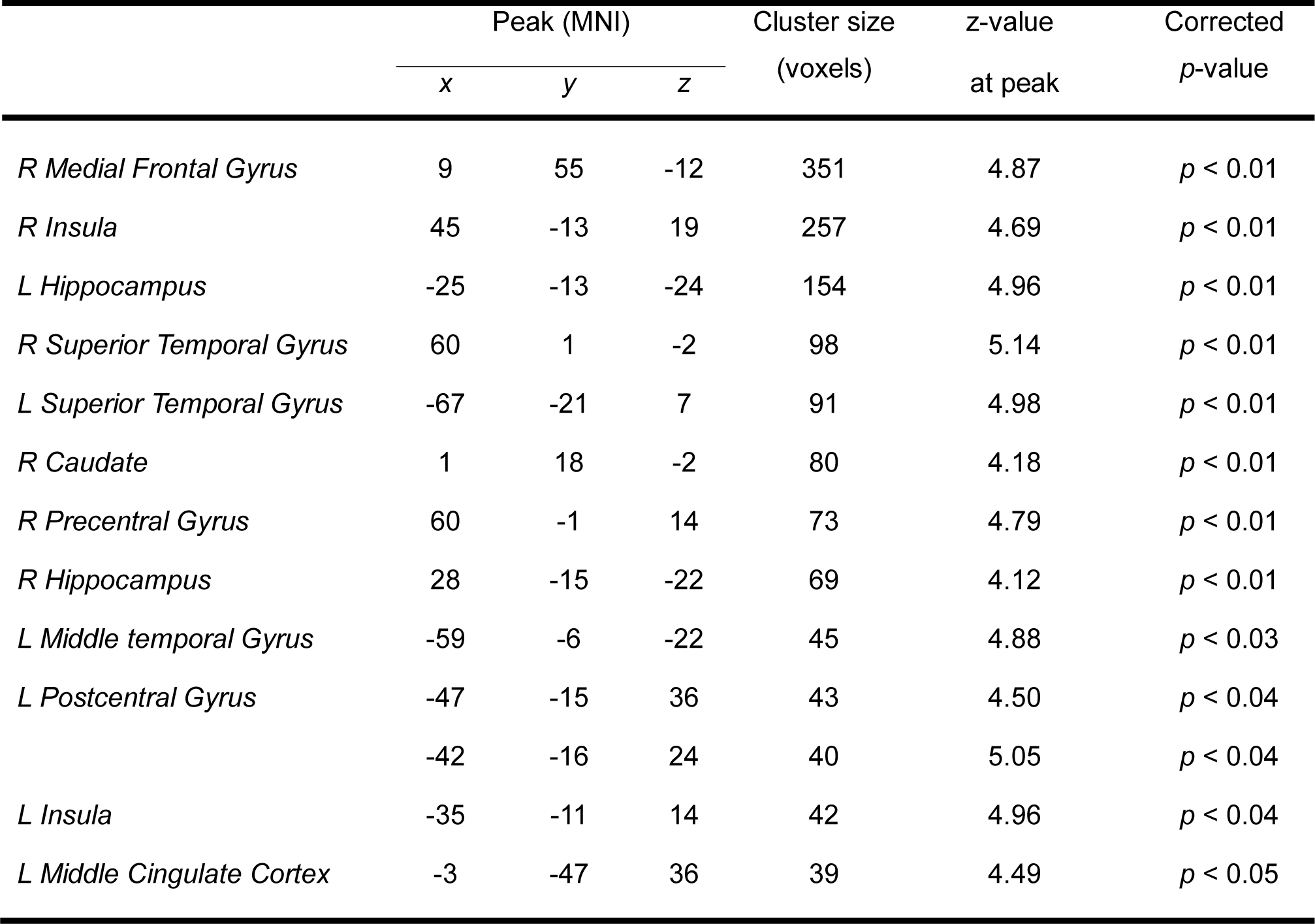
List of clusters significantly negatively interacting with motor errors

|  | Peak (MNI) |  |  | Cluster size<br>(voxels) | z-value<br>at peak | Corrected<br>*p*-value |
| --- | --- | --- | --- | --- | --- | --- |
|  | *x* | *y* | *z* |  |  |  |
| *R Medial Frontal Gyrus* | 9 | 55 | -12 | 351 | 4.87 | $p < 0.01$ |
| *R Insula* | 45 | -13 | 19 | 257 | 4.69 | $p < 0.01$ |
| *L Hippocampus* | -25 | -13 | -24 | 154 | 4.96 | $p < 0.01$ |
| *R Superior Temporal Gyrus* | 60 | 1 | -2 | 98 | 5.14 | $p < 0.01$ |
| *L Superior Temporal Gyrus* | -67 | -21 | 7 | 91 | 4.98 | $p < 0.01$ |
| *R Caudate* | 1 | 18 | -2 | 80 | 4.18 | $p < 0.01$ |
| *R Precentral Gyrus* | 60 | -1 | 14 | 73 | 4.79 | $p < 0.01$ |
| *R Hippocampus* | 28 | -15 | -22 | 69 | 4.12 | $p < 0.01$ |
| *L Middle temporal Gyrus* | -59 | -6 | -22 | 45 | 4.88 | $p < 0.03$ |
| *L Postcentral Gyrus* | -47 | -15 | 36 | 43 | 4.50 | $p < 0.04$ |
| | -42 | -16 | 24 | 40 | 5.05 | $p < 0.04$ |
| *L Insula* | -35 | -11 | 14 | 42 | 4.96 | $p < 0.04$ |
| *L Middle Cingulate Cortex* | -3 | -47 | 36 | 39 | 4.49 | $p < 0.05$ |

## 3. Results

### 3.1. Successful motor and episodic learning

Motor learning was successful, as indicated by decreases in motor error within feedback blocks (for all blocks, 1-tailed t-test: *df* = 23, *p* < 0.0017, Cohen’s *d* = 0.69) that plateaued within non-feedback blocks (Figure 2A). Performance worsened between the last trial of a feedback block and the first trial of the next feedback block of the same type due to forgetting, but increased overall across successive feedback and non-feedback blocks, as in prior studies ^4,33^. In order to prevent premature learning and reliance on habitual responding, each subject successfully learned two distinct visuomotor rotations (+40 and −40 degrees) ^4,33,34^. For each subject, errors in the last blocks for each rotation (feedback and non-feedback) were significantly lower than the imposed rotations (all *p* < 10^−7^). There was no difference in overall motor learning of the two rotations (*T*_(22)_ = 1.58, *p* = 0.13), and thus subsequent analysis collapsed these conditions. Control and non-feedback blocks were used as control conditions for which learning should have been minimal or absent. Indeed, there was no significant learning for most of control and non-feedback blocks (13 out of 16; *p* > 0.05, 1-tailed t-test) except for three non-feedback blocks in which marginal learning occurred (*p value range* = 0.022-0.048). Thus, motor learning was highly successful and based almost entirely on feedback.

Memory for object-location associations studied in an interleaved fashion during motor learning was assessed using a two-step test, whereby each trial was tested for recognition memory of the object followed immediately by recollection of its studied location. Recognition memory was successful, as indicated by accurate discrimination of studied objects from new/unstudied objects (Figure 2B). Subjects responded “old” with higher confidence to old objects and “new” with higher confidence to new objects (*F*_(3,66)_ = 125.0, *p* < 0.001, η_p_^2^ = 0.85 for the interaction of response type by object old/new status in a two-way repeated-measures ANOVA). All the subjects performed above chance, as indicated by discrimination sensitivity scores (*d*’) computed irrespective of confidence that were greater than zero (mean=1.36, range=0.30-2.59).

Recall of object locations was also successful and was strongly related to high-confidence object recognition ^35^. Object-location recall rates were significantly higher than the chance level of 0.25 (Figure 2C) when subjects recognized objects with high confidence, (0.66±0.040; *T*_(22)_ = 10.28, *p* < 0.001, Cohen’s *d* = 2.14) and low confidence (0.40±0.045; *T*_(21)_ = 3.45, *p* < 0.005, Cohen’s *d* = 0.74). Recall rates were not significantly higher than chance when subjects failed to recognize objects regardless of confidence (0.23±0.049; *T*_(17)_ = 0.41, *p* = 0.69) (Figure 2C). Object recognition and object-location recall were highly correlated across subjects (*r* = 0.92, *p* < 0.001), indicating that both were roughly equivalent indicators of strong episodic memory. In subsequent analysis, we primarily considered recognition memory confidence when assessing the interaction of motor with episodic learning, as this provided a graded episodic memory outcome.

### 3.2. Motor learning impaired episodic learning

To identify the influence of motor learning on episodic learning, a two-way repeated-measures ANOVA was performed on recognition memory performance, with subjects as random factors, and motor-feedback conditions (control, feedback, and non-feedback) and three experimental runs as repeated-measures. Episodic memory varied for the control, feedback, and non-feedback conditions irrespective of run (main effect *F*_(2,44)_ = 14.44, *p* < 10^−4^, η_p_^2^ = 0.40), and this effect of feedback conditions varied across runs (interaction effect *F*_(4,88)_ = 4.30, *p* < 0.005, η_p_^2^ = 0.16). Post-hoc pairwise tests indicated that recognition memory was worse for items learned in feedback blocks (when motor learning occurred) than in control blocks (*T*_(22)_ = 3.99, uncorrected *p* < 0.001, Cohen’s *d* = 0.85) and in non-feedback blocks (*T*_(22)_ = 5.29, uncorrected *p* < 0.001, Cohen’s *d* = 1.13), when no motor learning occurred (Figure 3A). We also tested whether nonspecific effect such as attention accounts for the difference in recognition memory between the conditions. Interestingly, reaction time varied for the three conditions, increasing in the order of control, feedback, and non-feedback conditions (*F*_(2,44)_ = 20.80, *p* < 0.001, η_p_^2^ = 0.49). Post-hoc pairwise tests indicated, in feedback blocks, the reaction time is shorter than in control blocks (*T*_(22)_ = 2.37, uncorrected *p* = 0.027), but longer than in non-feedback blocks (*T*_(22)_ = 4.30, uncorrected *p* < 0.001). However, when combining control and non-feedback conditions as a non-motor learning condition, reaction time is not different from that in a motor learning condition (feedback) (*T*_(22)_ = 0.94, *p* = 0.36). Thus, attention, which could be measured by reaction time, may not account for worse recognition memory in motor learning blocks. Furthermore, there was no significant difference in recognition memory for items learned in control blocks versus non-feedback blocks (*T*_(22)_ = 0.74, *p* = 0.46) despite of large difference in reaction time between two conditions (*T*_(22)_ = 6.08, *p* < 0.001), with performance for these conditions highly correlated across participants (*r* = 0.72, *p* < 0.001). In sum, motor learning that occurred during feedback blocks was associated with worse episodic learning, relative to non-feedback and control blocks. Non-feedback and control blocks were included to guard against nonspecific influences of motor and attentional demands on episodic memory ^36–38^, and they were roughly matched in any influence they had on episodic memory.

To determine whether the amount of motor learning correlated negatively with the success of episodic learning, we computed the average motor learning within individual learning blocks and the success of recognition memory for the same blocks. These values were highly negatively correlated (*r* = 0.76, *p* < 0.001, Figure 3B), indicating that the relative success of motor learning was highly related to the relative failure of episodic learning.

### 3.3. Episodic learning impaired motor learning

Episodic memory formation impaired the motor learning that occurred on the trials following the object-location encoding. Trial-by-trial motor learning was calculated as the improvement (decrease of error) from the current to the next trial. We assessed this improvement value following object-location trials that were categorized into four increasing levels of episodic learning success based on subsequent performance (later-missed, later-hit with low-confidence, later-hit with high-confidence, and later-hit with high-confidence and source-correct). Motor improvement scores varied by episodic learning success (main effect *F*_(3,22)_ = 48.5, *p* < 0.001, η_p_^2^ = 0.87), with less improvement occurring for relatively more successful episodic learning (Figure 3C). Post-hoc pairwise tests indicated less motor improvement for the highest level of episodic learning success compared to all lower levels (*p* < 0.008; Figure 3C) and for the next-to-highest level of episodic learning success compared to all lower levels (*p* < 0.02; Figure 3C), with no significant difference in motor improvement for the two lowest levels of episodic learning success (*T*_(22)_ = 1.32, *p* = 0.20). Thus, just as there was a negative impact of motor learning on the immediately forthcoming episodic learning trial, episodic learning disrupted the forthcoming motor learning.

### 3.4. Network activity reflecting episodic learning impairment by motor learning

To identify brain activity reflecting the negative impact of motor learning on episodic learning, we first modeled fMRI activity reflecting trial-by-trial changes in motor performance error, which reflects the magnitude of motor learning ^2–4,13,17,28–31^. Activity of a hippocampal-prefrontal network was negatively modulated by motor learning, such that activity in these areas, which are strongly associated with episodic memory ^39–41^, was lower when more motor learning occurred on the immediately preceding trial (Figure 4A, Table 1). Furthermore, subjects with greater levels of episodic learning impairment by motor learning (difference in recognition memory for items studied during feedback versus non-feedback blocks) also had greater negative modulation of fMRI activity by motor learning within the vmPFC (*r* = 0.53, *p* = 0.01; Figure 4B), which was the focus of the analysis as it was the location with the most robust negative modulation of fMRI activity by motor learning (Table 1). When we controlled out the potential effect of reaction time on the modulation given significant difference in reaction time between feedback and non-feedback blocks (*T*_(22)_ = 4.30, uncorrected *p* < 0.001), the relationship remained significant (*r* = 0.54, *p* = 0.01). These findings indicate that successful motor learning impaired subsequent episodic learning and decreased fMRI signals of episodic learning in the episodic network.

**Figure 4.**
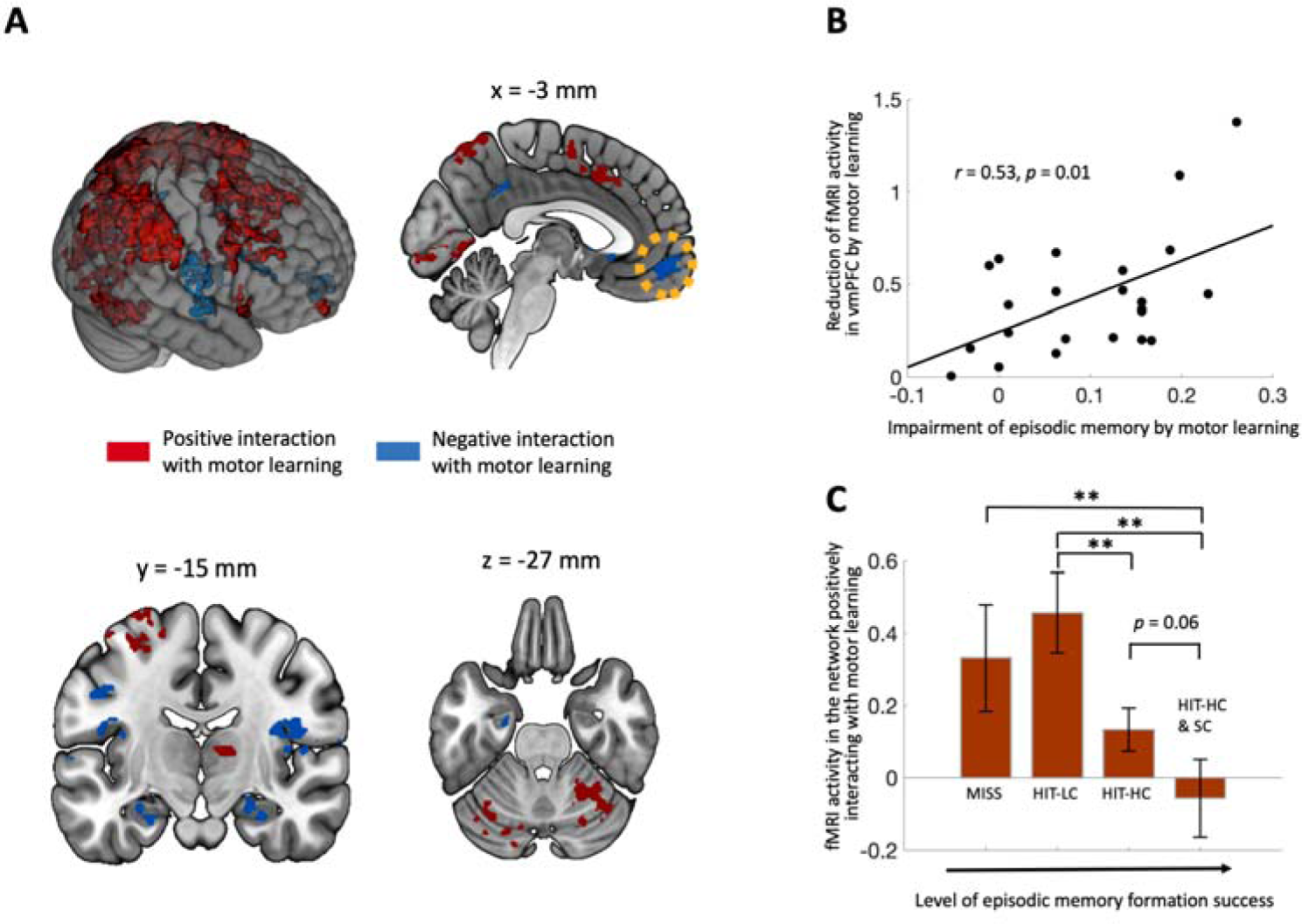
Reciprocal interference of brain activity reflecting motor and episodic learning. (**A**) Motor learning correlated positively (red) with activity of a prototypical prefronto-parieto-cerebellar motor-learning network and negatively (blue) with activity of a prototypical hippocampal-prefrontal episodic network. (**B**) The extent of negative modulation of fMRI activity within vmPFC, marked with a yellow-dotted circle in (A), predicted the level of episodic learning impairment by motor learning shown in Figure 3A. (**C**) fMRI activity of the motor learning network (positive clusters in A) decreased with higher level of episodic memory formation success.

### 3.5. Network activity reflecting motor learning impairment by episodic learning

Activity within a prototypical prefronto-parieto-cerebellar network associated with motor learning ^4,42–45^ was positively modulated by trial-by-trial motor performance error (Figure 4A, Table 2). We tested whether increasing levels of episodic learning success was associated with subsequent reductions in fMRI correlates of motor learning within this network, which would mirror the effects identified on motor learning behavioral performance (Figure 3C). Indeed, activity related to motor learning varied by the four levels of episodic learning success (main effect *F*_(3,22)_ =6.30, *p* < 0.001, η_p_^2^ = 0.46) (Figure 4C), with post-hoc pairwise tests indicating that this was due to decreased activity with higher levels of episodic learning success (Figure 4C), as was the case for the effects on behavior. This relationship was specific to the motor-learning network, with no similar relationship in the regions that negatively interacted with motor learning (*F*_(3,22)_ = 1.89, *p* = 0.14). Thus, successful episodic memory formation was associated with worse subsequent motor learning and reduced fMRI signals of motor learning in the prefronto-parieto-cerebellar motor network.

**Table 2.** List of clusters significantly positively interacting with motor errors

|  | Peak (MNI) |  |  | Cluster size<br>(voxels) | z-value<br>at peak | Corrected<br>p-value |
| --- | --- | --- | --- | --- | --- | --- |
|  | x | y | z |  |  |  |
| *R Supramarginal Gyrus* | 59 | -28 | 46 | 5475 | 5.56 | $p < 0.01$ |
| *L Inferior Parietal Lobule* | -59 | -35 | 48 | 3455 | 5.48 | $p < 0.01$ |
| *R Middle Frontal Gyrus* | 40 | 24 | 41 | 856 | 5.05 | $p < 0.01$ |
| | 35 | 65 | -10 | 39 | 4.12 | $p < 0.01$ |
| *R Cuneus* | 16 | -79 | 9 | 789 | 4.89 | $p < 0.01$ |
| *R Middle Cingulate Cortex* | 4 | 18 | 44 | 719 | 5.28 | $p < 0.01$ |
| *R Cerebellum (Lobule VI)* | 23 | -50 | -29 | 633 | 5.12 | $p < 0.01$ |
| *R Superior Frontal Gyrus* | 23 | 1 | 68 | 605 | 5.49 | $p < 0.01$ |
| *R Cerebellum (Lobule VI)* | -31 | -66 | -25 | 293 | 4.69 | $p < 0.01$ |
| *R Middle Occipital Gyrus* | 40 | -81 | 34 | 275 | 4.85 | $p < 0.01$ |
| *R Inferior Frontal Gyrus* | 47 | 26 | 29 | 148 | 4.74 | $p < 0.01$ |
| | 57 | 18 | 5 | 115 | 4.49 | $p < 0.01$ |
| *L Cerebellum (Crus 1)* | -16 | -83 | -27 | 112 | 4.71 | $p < 0.01$ |
| *R Middle Temporal Gyrus* | 65 | -57 | -3 | 94 | 4.19 | $p < 0.01$ |
| *L Supplementary Motor Area* | -3 | -4 | 60 | 79 | 4.98 | $p < 0.01$ |
| *L Superior Parietal Lobule* | -33 | -52 | 70 | 52 | 4.28 | $p < 0.02$ |
| *R Thalamus* | 11 | -15 | 9 | 45 | 5.18 | $p < 0.03$ |
| *L Middle Frontal Gyrus* | -48 | 24 | 32 | 42 | 4.40 | $p < 0.04$ |
| | -42 | 35 | 32 | 38 | 3.82 | $p < 0.05$ |

Finally, given that we found behavioral and fMRI evidences supporting bidirectional interference between episodic and motor learning networks, we expected to identify signatures of functional coupling between the learning networks from the PPI analyses. However, there were no regions in the whole brain showing significant PPI (i.e., modulation of functional connectivity between the networks by the degree of episodic or motor learning).

## 4. Discussion

A novel fMRI experiment allowed measurement of trial-by-trial motor learning and the corresponding level of episodic memory formation success, providing behavioral and fMRI evidence supporting bidirectional interference between these systems. Previous findings have shown competition between motor versus episodic memory systems for common learning goals. The competition varies across different stages of motor learning, producing automaticity ^7^ with decreasing dependence on episodic memory ^11^. Motor and episodic memory systems also competitively interact during consolidation given distinct learning goals specific to each system ^12–14^. The current results demonstrate these memory systems also compete during memory formation even when system-specific information is presented at distinct times. Thus, we provide another line of evidences supporting competitive interaction between two memory systems traditionally viewed as encapsulated and separated ^1,46^.

Competitive interactions such as those identified here have also been seen for other types of learning systems. For instance, in reward-based learning involving competition between hippocampus and striatum ^47,48^. However, like previous studies investigating motor-episodic competition during motor learning, studies of reward-based learning have typically observed hippocampal-striatal competition when subjects are tasked with learning one piece of information that could be relevant to both systems. Here, we segregated to-be-learned information into discrete yet temporally proximal packets, and nonetheless identified competition.

The source of the competitive interaction is unclear and could be due to increased demands such as task switching ^49–51^. Our finding that greater vmPFC deactivation was related to more interference suggest competition in some aspect of information integration functions supported by this region ^52–55^. Most studies employing a dual-task paradigm aimed to understand interference presented in performing tasks *per se* by measuring reaction time and found overlapped neural representation between two tasks, supporting interference due to competition for the same brain region ^36,56^. In contrast, our findings support separate neural substrates of different memory systems and attenuated activation in each system as a neural signature of the interference by the other memory processing although PPI analyses failed to identify direct functional coupling between memory systems. However, it is also possible that two learning systems are partially overlapped for common memory resource. Specifically, hippocampus processes not only episodic information but also motor information ^57,58^ and cerebellum, in addition to its motor functions, is also engaged in a variety of cognitive functions including episodic memory processing ^59–61^. For the future works, like a previous study ^14^, brain stimulation targeting vmPFC with functional connectivity analysis could be considered to further investigate a potential role of the region in arbitrating motor-episodic competition.

One limitation of our study is that the design did not permit segregation of motor learning into fast/declarative versus slow/adaptation components ^8,11,13^. Previous studies have identified episodic memory tasks interfered with motor learning during consolidation particularly for the fast/declarative component ^13^. However, we have not tested this hypothesis in the current experiment. For this, we should measure delayed retention of motor learning, which is predictive of the slow component ^62^, and design separate control trials without interfering episodic memory task (e.g., showing noise instead of objects) to test whether episodic learning affects the delayed retention of motor memory. For the other direction of interference, motor memory to episodic memory, either of fast or slow component seems to disrupt episodic memory given that the effect tended to decrease but remained significant in late stages of learning, when slow/adaptation components dominate performance. Nonetheless, stronger evidence could be obtained in future studies by segregating these components of motor learning and testing for their selective interaction with episodic learning.

Another potential limitation of our findings is that it is not fully clear whether the interference effects were anterograde versus retrograde. This is because episodic and motor learning events were interleaved and consecutive. Future studies could address this question by varying the time gaps between episodic and motor learning demands ^18^ and/or by varying their order across trials.

In summary, our results provided behavioral and fMRI evidence supporting competitive reciprocal interaction between motor and episodic learning despite system-specific learning goals. Additional research is needed to further evaluate mechanisms for such competition and to test whether either of the systems is particularly dominant for producing such interaction, such as by selectively modulating one system versus the other using network-targeted brain stimulation ^14,15,19,63,64^.

## 6. Data Availability

All MRI data used in this study are archived in the Northwestern University Neuroimaging Data Archive (NUNDA, https://nunda.northwestern.edu) and can be downloaded following registration. The corresponding author can provide any information on dataset identification within the NUNDA system if necessary. Additional data related to this paper may be requested from the corresponding author.

## 8. Acknowledgements

We thank Jonathan O’Neil for assistance with data collection. Neuroimaging was performed at the Northwestern University Center for Translational Imaging, supported by the Northwestern University Department of Radiology. This research was also supported in part through the computational resources and staff contributions provided for Quest, the high-performance computing facility at Northwestern University.

## 9. Author contributions

S.K. designed the study, performed the experiment, analyzed data and wrote the manuscript.

